# Membrane-assisted Aβ40 aggregation pathways

**DOI:** 10.1101/2024.09.05.611426

**Authors:** Fidha Nazreen Kunnath Muhammedkutty, Huan-Xiang Zhou

**Affiliations:** Department of Chemistry, University of Illinois Chicago, Chicago, IL, USA; Department of Physics, University of Illinois Chicago, Chicago, IL, USA

## Abstract

Alzheimer’s disease (AD) is caused by the assembly of amyloid-beta (Aβ) peptides into oligomers and fibrils. Endogenous Aβ aggregation may be assisted by cell membranes, which can accelerate the nucleation step enormously, but knowledge of membrane-assisted aggregation is still very limited. Here we used extensive MD simulations to structurally and energetically characterize key intermediates along the membrane-assisted aggregation pathways of Aβ40. Reinforcing experimental observations, the simulations reveal unique roles of GM1 ganglioside and cholesterol in stabilizing membrane-embedded β-sheets and of Y10 and K28 in the ordered release of a small oligomeric seed into solution. The same seed leads to either an open-shaped or R-shaped fibril, with significant stabilization provided by inter- or intra-subunit interfaces between a straight β- sheet (residues Q15-D23) and a bent β-sheet (residues A30-V36). This work presents the first comprehensive picture of membrane-assisted aggregation of Aβ40, with broad implications for developing AD therapies and rationalizing disease-specific polymorphisms of amyloidogenic proteins.

## Introduction

The aggregation of amyloid beta (Aβ) peptides into oligomers and fibrils is a key feature of Alzheimer’s disease (AD). These peptides are produced in a membrane environment upon the cleavage of the amyloid precursor protein by the sequential action of β and γ secretases at an extracellular site and a transmembrane site, to generate the N- and C-termini, respectively, of Aβ40/42 ^1, 2^, with K28 serving as a membrane anchor ^3^ (Fig. S1A). The C-terminal portion (A30 to the C-terminus) of Aβ40/42, being completely nonpolar, is a part of the transmembrane helix of the precursor protein ^4^ and thus expected to be favored in a membrane environment. It has been proposed that a substantial amount of Aβ peptides is associated with cell membranes ^5, 6^. Indeed, Aβ peptides bind to cell membranes in an oligomeric state ^7^, and these membrane-bound oligomers can cause cytotoxicity by disrupting membranes via forming ion channels ^8, 9^, creating defects ^10^, or extracting lipids ^11, 12^.

Lipid membranes or membrane mimetics such as detergent micelles can significantly promote the formation of β-sheet-rich aggregates for Aβ peptides ^13^. Roles of specific lipids including monosialotetrahexosylganglioside (GM1) and cholesterol in the membrane-assisted aggregation of Aβ peptides have been recognized ^14, 15^. Residues important for membrane interactions have also been identified, including Y10 and K28 ^9, 16^. In several solid-state nuclear magnetic resonance (NMR) and cryo-electron microscopy (cryo-EM) studies, the fibril structures resulting from membrane-assisted aggregation of Aβ40 have been characterized, including one formed on exposure to DMPC/DMPG vesicles ^17^, one via association with rat synaptic plasma membranes ^18^, and six polymorphic structures formed in the presence of DMPG vesicles ^19^. However, there is only limited knowledge of the membrane-assisted aggregation pathways or the intermediates formed. In this regard, the work of Kenyaga et al. ^18^ is particularly valuable. By focusing on intermediates formed in the lag phase of fibrillation, these authors developed a schematic mechanism for the membrane-associated nucleation step, identifying the assembly of the C-terminal β-sheet as a key event. A recent study using solid-state NMR and molecular dynamics (MD) simulations defined a detergent-assisted oligomerization pathway for Aβ42, where SDS and DPC micelles stabilize an antiparallel β-sheet tetramer that can further aggregate into a high molecular weight (∼150 kDa) oligomer ^20^. The latter oligomer has been structurally characterized by solid-state NMR and cryo-EM and can potentially act as a transmembrane ion channel ^21^.

Over thirty Aβ40 fibril structures have been reported. A common characteristic is the presence of two β-sheets; one, to be termed N-sheet, is formed by residues Q15-D23, and the other, to be termed the C-sheet, is formed by residues A30-V36 (Fig. S1B). Eight *in vitro* fibril structures consist of straight N- and C-sheets (Fig. S1C) ^22–26^. Other fibril structures, with three exceptions, were formed in lipid environments or derived from patient brains, and all have a straight N-sheet but a C-sheet with a sharp bent at G33. Frieg et al. ^19^ solved six lipidic fibril structures. The most populated one is dimeric, with the N- and C-sheets of each fibrillar subunit adopting an open shape and the two subunits forming two identical interfaces, each between a straight N-sheet and the convex side of a bent C-sheet (Fig. S1D). The five less populated lipidic fibrils are monomeric, where the C-sheet curves onto the N-sheet (Fig. S1E). Seven patient-derived and two *in vitro* fibrils also have the open-shaped subunit of Fig. S1D ^27–31^, while another patient-derived fibril has the curved subunit of Fig. S1E ^32^. The membrane-exposed fibrils resolved by Niu et al. ^17^ and by Kenyaga et al. ^18^ have an R-shaped subunit, with an intra-subunit interface between the straight N-sheet and the convex side of the bent C-sheet, in a monomeric form and a dimeric form (Fig. S1F), respectively. In the latter case, the inter-subunit interface involves the concave side of the bent C-sheet. Four human or mouse brain-derived fibril structures ^33, 34^ and one *in vitro* structure ^35^ also have an R-shaped subunit, in either dimeric or tetrameric form (Fig. S2).

Patient-derived fibrils of amyloidogenic proteins such as tau, α-synuclein, and TDP-43, in addition to Aβ, are found to exhibit disease-specific structures. Tau, for instance, forms distinct fibril folds in different neurodegenerative conditions like AD ^36, 37^, Pick’s disease ^38^, chronic traumatic encephalopathy ^39^, corticobasal degeneration ^40^, and progressive supranuclear palsy ^41^. Similarly, TDP-43 shows unique fibril structures in amyotrophic lateral sclerosis ^42^ and frontotemporal lobar degeneration ^43^, while α-synuclein exhibits structural polymorphism in multiple system atrophy ^44^ and Parkinson’s disease ^45^. Aβ42 forms distinct fibril folds in familial and sporadic AD ^46^. For Aβ40, distinct fibril structures were derived from patients with different AD-related diagnoses. Those from sporadic AD ^28, 31^, sporadic or Dutch-type cerebral amyloid angiopathy ^30^, and sporadic AD and cerebral amyloid angiopathy ^27, 29^ have the open-shaped fibrillar subunit of Fig. S1D; one from a mixed diagnosis of AD and Lewy body dementia ^32^ has the curved fibrillar subunit of Fig. S1E; and those from AD patients with Down syndrome ^33^ or the Arctic mutation (E22G) ^34^ have the R-shaped subunit of Fig. S1F. Disease-specific fibril structures are likely dictated by aggregation pathways, e.g., due to environmental factors associated with particular disease conditions. Therefore studying aggregation pathways is crucial for understanding disease mechanisms and developing targeted therapies.

MD simulations can potentially generate atomic-level information on membrane-assisted aggregation that is not amenable to experimental techniques. Monomeric as well small oligomeric Aβ peptides were simulated in membrane environments, revealing specific lipid-peptide interactions ^15, 47–50^. For example, GM1 ganglioside has high affinity for Aβ42 dimers ^48^; cholesterol enhances the membrane binding of Aβ42 monomers and dimers and promotes structural changes, in particular β-sheet formation ^15^; certain residues including Y10 and K28 mediate membrane binding of Aβ42 monomers and dimers ^48, 50^. For other amyloidogenic proteins, MD simulations have demonstrated membrane-assisted oligomerization of amylin, where monomers initially bind to membranes in an α-helical conformation and then self-assemble into a β-sheet through intermonomer interactions ^51^. Simulations of the FUS low-complexity fibrils at membrane surfaces suggest stabilization due to strengthened backbone hydrogen bonds ^52^. Still, for Aβ peptides and many other amyloidogenic proteins, how membranes assist their aggregation remains poorly defined.

In this study, we used conventional, steered, and targeted MD simulations, in addition to umbrella sampling, to structurally and energetically characterize key intermediates along the membrane-assisted aggregation pathways of Aβ40. Reinforcing experimental observations, the simulations reveal unique roles of GM1 and cholesterol in stabilizing membrane-embedded β-sheets and of Y10 and K28 in the ordered release of a small oligomeric seed into the aqueous solution. Moreover, the same seed leads to either the open-shaped or the R-shaped fibril, with significant stabilization provided by inter- or intra-subunit interfaces between the straight N-sheet and the bent C-sheet. This work presents the first comprehensive picture of membrane-assisted aggregation of Aβ40, with broad implications for developing AD therapies and rationalizing disease-specific polymorphisms of amyloidogenic proteins.

## Results

Our aim was to map out the structural and energetic details of the major intermediates along the membrane-assisted aggregation pathways of Aβ40 using MD simulations. This effort was guided by a rough sketch of the aggregation process, including the assembly of a β-sheet-rich small oligomeric seed in the membrane environment, reconfiguration of the seed in the aqueous environment, and growth into fibrils.

### N- and C-sheets have disparate burial depths and stability in membranes

Membrane-assisted nucleation involves the insertion of Aβ40 monomers into membranes and the assembly of a small oligomeric seed ^18^. During this stage lasting several hours, the secondary structures change from initial partial helices to more unstructured to increasing β-sheets. Given that all known Aβ40 fibril structures consist of an N-sheet formed by residues Q15-D23 and a C-sheet formed by residues A30-V36 (Fig S1B), we hypothesized that the oligomeric seed contains one or both of these β-sheets. Our MD simulations focused on one important aspect: how are these β-sheets accommodated in the membrane environment? To address this question, we placed an N-sheet (four strands, each of the Q15-D23 fragment) and a C-sheet (four strands, each of the K28- G38 fragment) in membranes. The initial structures of these fragments were taken from the predominant fibril structure, Protein Data Bank (PDB) entry 6W0O, of AD patients ^28^. Our membranes mimicked the lipid composition of rat synaptic plasma membranes, with POPC, cholesterol, palmitoylsphingomyelin (PSM), POPE, POPI, GM1, and POPS at 46%, 25%, 12%, 7%, 5%, 3%, and 2%, respectively. The first four lipids are neutral whereas the last three are anionic.

To find a preferred burial depth in membranes, we placed each β-sheet at four different positions relative to the phosphate plane: one just beneath it, one further below, one just above it and within the headgroup region, and one just outside the membrane (Figs. 1A, B and S3). While the N-sheet started outside the membrane stays outside, it converges to a position intersecting the phosphate plane in the other three simulations within 100 ns (Fig. S3C). Likewise, the C-sheet started outside the membrane stays outside, but it converges to a position beneath the phosphate plane (Fig. S3D). The converged structures are shown in Fig. 1C, D. The deeper burial of the C-sheet relative to the N-sheet can be attributed to the fact that all the seven residues making up the C-sheet are nonpolar whereas four of the nine residues making up the N-sheet are polar (Fig. S1B). The deeper burial of the C-sheet is also confirmed by the placement (Fig. S4) predicted by the PPM web server ^53^, which is based on the transfer free energies of different amino acids from an aqueous solution to the membrane interface.

**Fig. 1.**
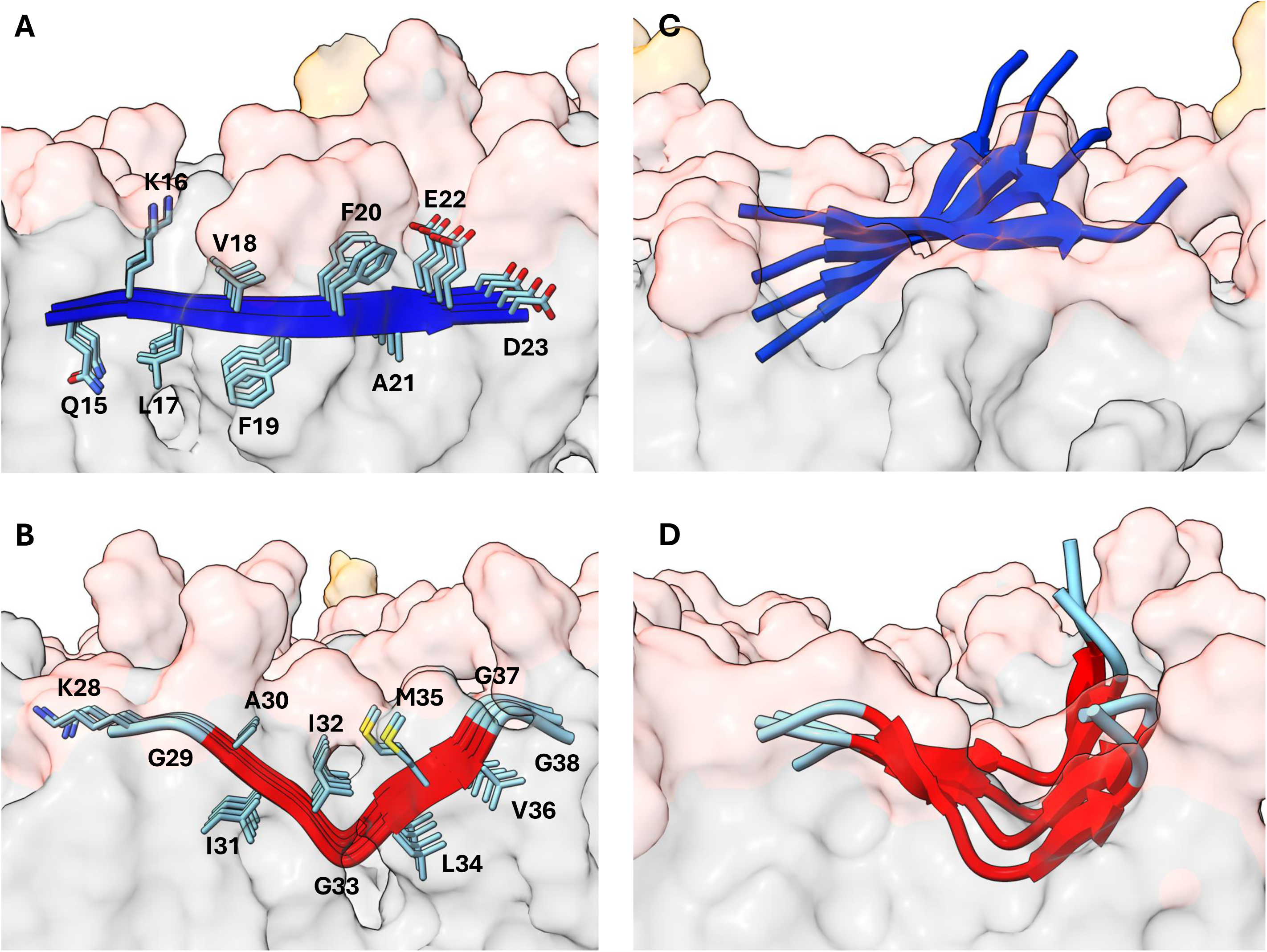
Disparate burial depths of the N- and C-sheets in membranes. (A, B) The N- and C-sheets at the deepest of four initial burial depths. (C) A snapshot of the N-sheet at 33.5 ns, just before the fraying of the β-sheet at the termini. (D) A snapshot of the C-sheet at 96.5 ns. The membrane is shown as surface, with the phosphate and all atoms above it in pink whereas those below the phosphate in gray. The hydroxyl group of cholesterol is in pink; the entire headgroup of GM1 is in orange.

In the converged structures (Figs. 1C, D and S3C, D), while the C-sheet has its two termini even with respect to the phosphate plane, the N-sheet has a distinct tilt with the C-terminus elevated relative to the N-terminus, due to electrostatic repulsion of the C-terminal E22 and D23 with anionic lipids. Both the N-sheet and each half of the bent C-sheet exhibit an expected ∼15° twist ^54^ (Fig. S5).

The secondary structure of the C-sheet is well maintained in the membrane, but the N-sheet shows fraying at the termini (Figs. 1C, D and S6). The higher stability of the C-sheet is partly consistent with the observation of Kenyaga et al. ^18^ that the C-sheet is formed in the membrane-assisted nucleation stage.

### Facile sidechain-lipid interactions stabilize Aβ40 tetramer in membranes

Next, we asked how the membranes stabilize an oligomeric seed. Using the converged structures of the N- and C-sheets in the preceding step as templates, we built a tetramer consisting of residues Y10-V40 in each chain. Due to the tilt of the N-sheet and the bend of the C-sheet, the intervening residues along with the C-terminus of the N-sheet form an arch; we will refer to the overall shape of this tetramer as “semi-open”. When placed in membranes, the tip of the arch is above the headgroups of all lipids except GM1 (Fig. 2A). In four replicate simulations lasting 600 ns each, the secondary structures of the semi-open tetramer are well maintained (Fig. S7). Again, the β- sheets exhibit a ∼15° twist (Fig. S5).

**Fig. 2.**
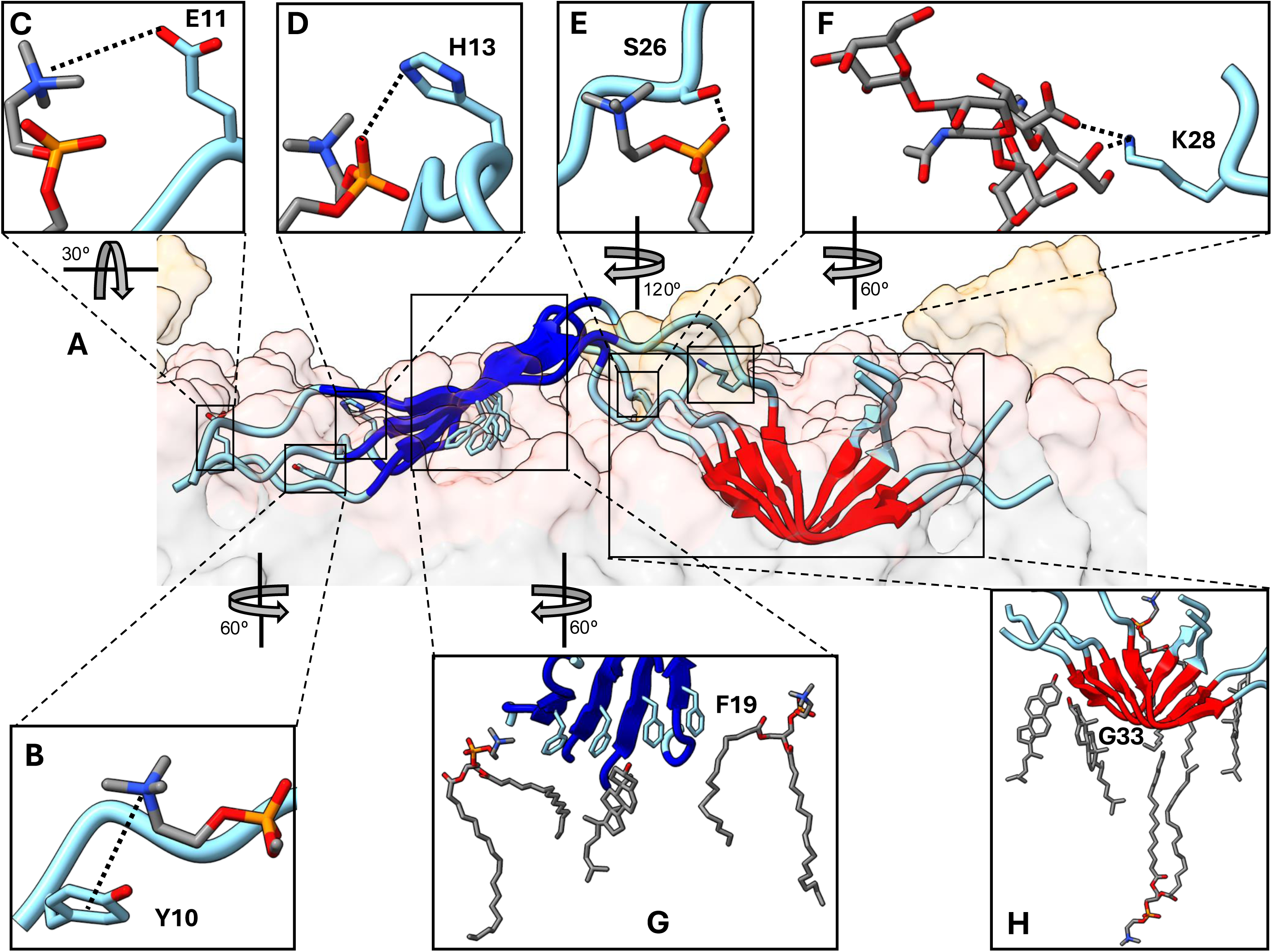
Interactions stabilizing the oligomeric seed in membranes. (A) A snapshot at 455.7 ns of one replicate simulation. The membrane has the same representation as in Fig. 1. The N- and C-sheets are in blue and red, respectively. (B-H) Zoomed views of peptide-lipid interactions. Salt bridges, hydrogen bonds, and cation-ρε interactions are indicated by thick dashed lines. Peptide carbon, lipid carbon, nitrogen, oxygen, and phosphorous are in cyan, grey, blue, red, and orange, respectively. Lipids are POPC, except for GM1 in (F); cholesterol is also present in (G) and (H).

Residues Y10, E11, H13, S26, and K28 in disordered regions and F19 and G33 within the β-sheets show high membrane-contact probabilities (Fig. 3A). Frequently, Y10 forms a cation-π interaction with a lipid choline, E11 forms a salt bridge with a choline, H13 and S26 form hydrogen bonds with lipid phosphates, and K28 forms salt bridges and hydrogen bonds with a GM1 headgroup (Fig. 2B-F).

**Fig 3.**
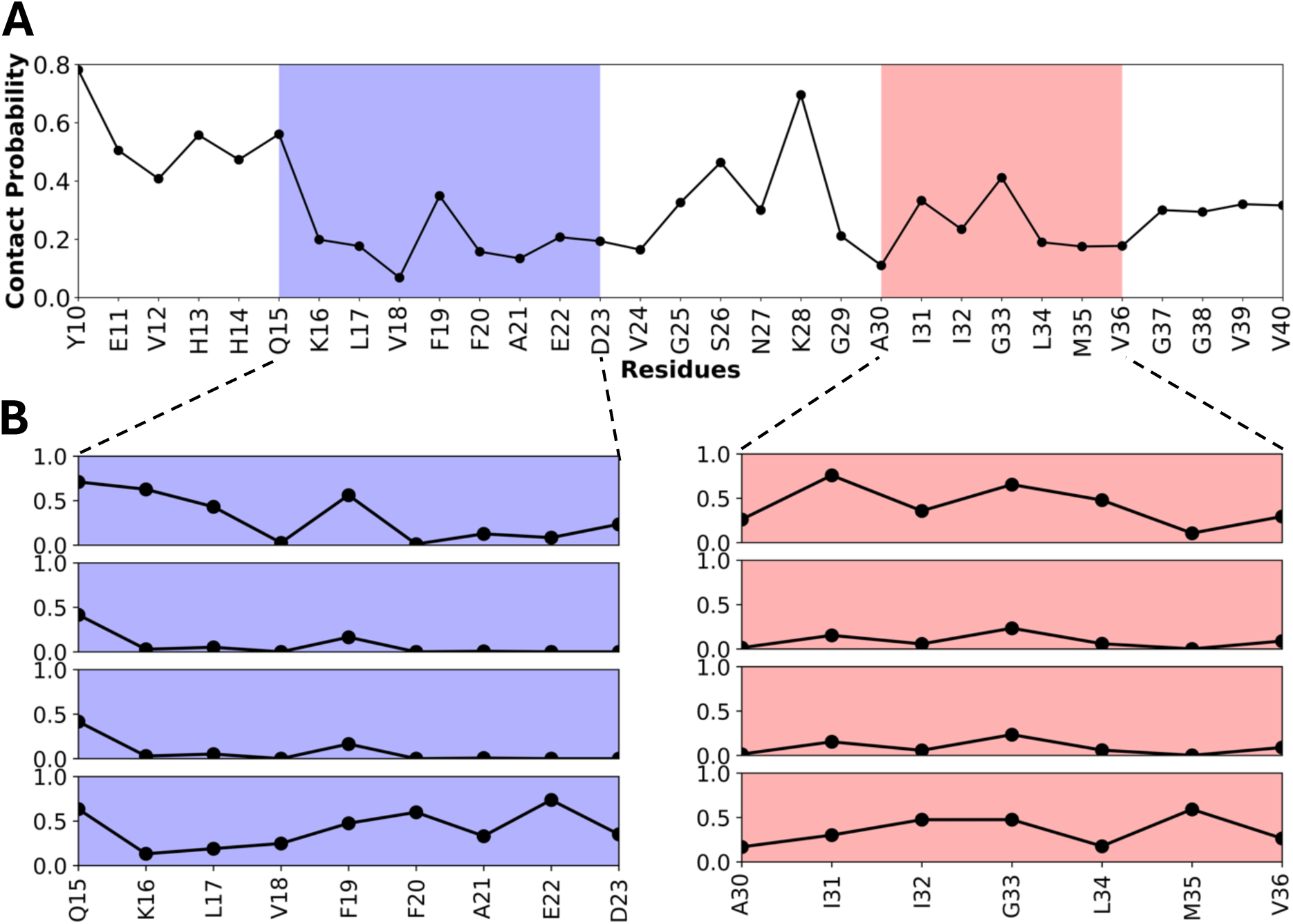
Membrane contact probabilities of individual residues in the oligomeric seed. (A) Membrane contact probabilities averaged over the four chains. (B) Corresponding results for the individual chains. Averages were taken over the entire trajectories of four replicate simulations. The N- and C-sheets are indicated by blue and red shading, respectively.

In the β-sheet regions, the F19 sidechain is the most deeply buried among the stretch of five nonpolar residues (L17-A21) in the N-sheet (Fig. 2G), while G33 is the most deeply buried residue in the C-sheet due to the sharp bent (Fig. 2H). Among the four strands of the N- or C-sheet, the two edge strands have higher membrane-contact probabilities than the two interior strands (Fig. 3B). This difference arises because the acyl chains of lipids (except cholesterol) bend to reach the underside of the β-sheet (Fig. 2G), thereby limiting access to the interior strands. For the C-sheet, due to its deep burial, acyl chains from the opposite leaflet can also reach the underside (Fig. 2H).

To ascertain the roles of different lipids in stabilizing the tetramer, we decomposed the membrane-contact probability of each residue according to lipid types (Fig. S8). POPC is the most frequent partner for residues throughout the amino-acid sequence, mainly because it is present at the highest fraction of all lipid types. Other lipids show region-specific preferences. In particular, GM1 shows a disproportionate preference for the residues between the N- and C-sheets, because its long and anionic headgroup is in a position to interact with the elevated K28 via salt bridges and hydrogen bonds (Fig. 2F). Cholesterol, on the other hand, shows preference for the undersides of the N- and C-sheets, as the absence of a headgroup allows it to get under the β-sheets (Fig. 2G, H).

### Y10 tethers the N-sheet to membranes while the C-sheet is released

After the small oligomeric seed is formed in the membranes, it is released into the aqueous solution. The release is slow and may require the transient generation of membrane defects. Here we used steered MD simulations to accelerate this process. In four replicate simulations, the center of mass (COM) of all C_*α*_ atoms in the semi-open tetramer was pulled at a constant speed of 1 Å/ns away from the membrane (Fig. 4A). After 12-16 ns of simulations, the C-sheet is about to be released from the membrane and the membrane tether by K28-phosphate (or carboxyl) salt bridges is about to break, while the N-sheet is still tethered to the membrane by choline-Y10 cation-ρε interactions (Fig. 4B).

**Fig. 4.**
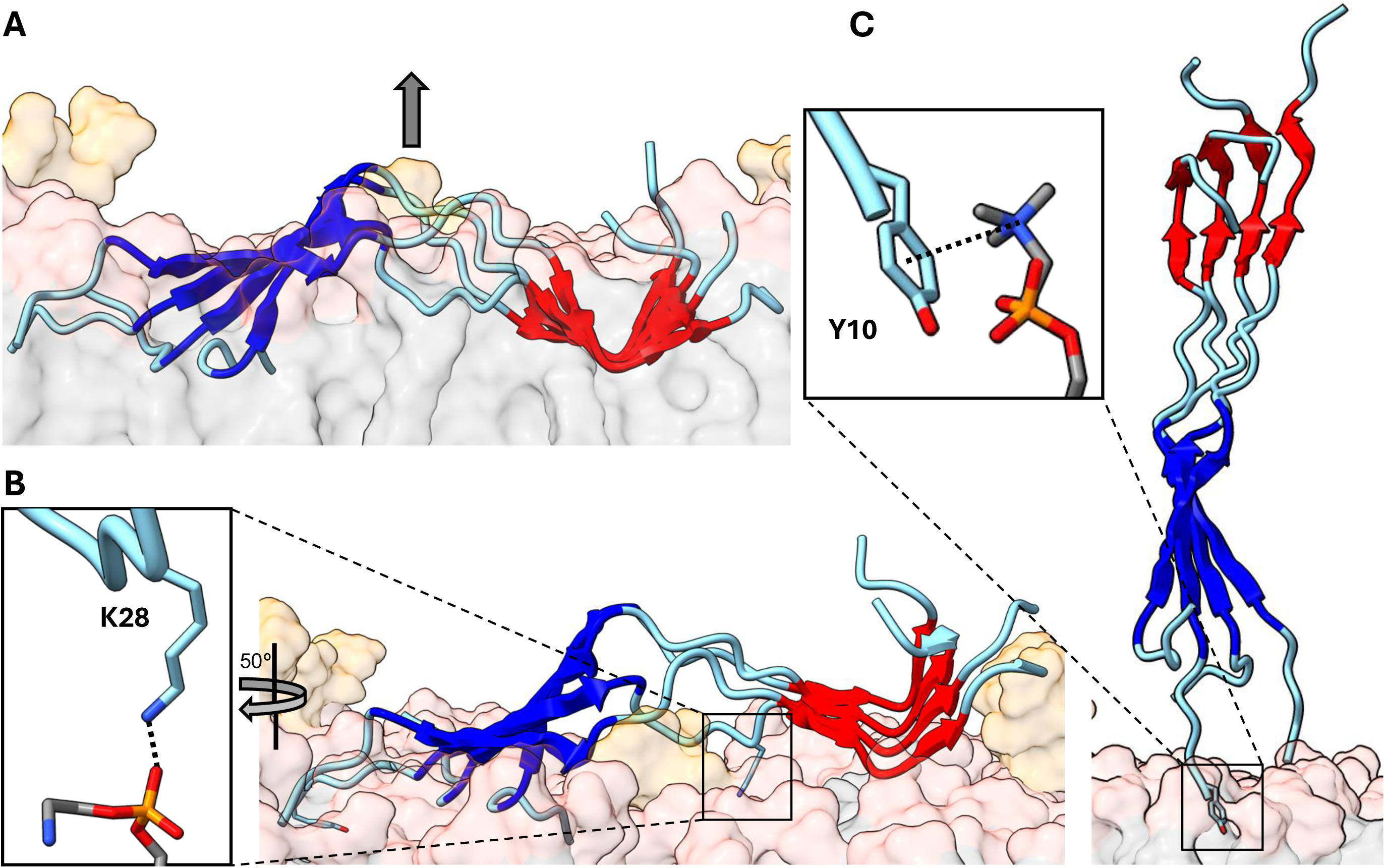
Release of the oligomeric seed from membrane to solution. (A) Initial snapshot of the steered MD simulations. The direction of pulling is indicated in an arrow. (B) A snapshot at 12.2 ns of one replicate simulation, where the C-sheet is almost entirely freed from the membrane and the K28 tether (zoomed view) is about to break. (C) A snapshot at 48.0 ns, where the entire oligomer is freed from the membrane, except for the Y10 tether (zoomed view). The oligomer adopts an open shape due to the pulling.

By ∼30 ns of simulations, the N-sheet is also freed from the membrane, except for the very N-terminus. The shape of the tetramer now changes from semi-open to open, with the arch straightened by the pulling force. The latter shape is maintained (Fig. 4C) till the choline-Y10 tether is finally broken around ∼50 ns.

### The interface between N- and C-sheets dominates fibril stability but that between C-sheets provides additional stability

After its release into the aqueous solution, the small oligomeric seed evolves into protofilaments for fibril growth. Specifically, the oligomeric seed, after an appropriate shape change, becomes a fibrillar subunit, which can dimerize to form a protofilament. We used targeted MD, steered MD, and umbrella sampling to map out the pathways and free-energy differences for the conversion of the fibrillar subunit among different shapes and the assembly of differently-shaped dimeric protofilaments (Fig. 5A-E). Here the peptide constructs are composed of chains comprising residues Q15-G38.

**Fig. 5.**
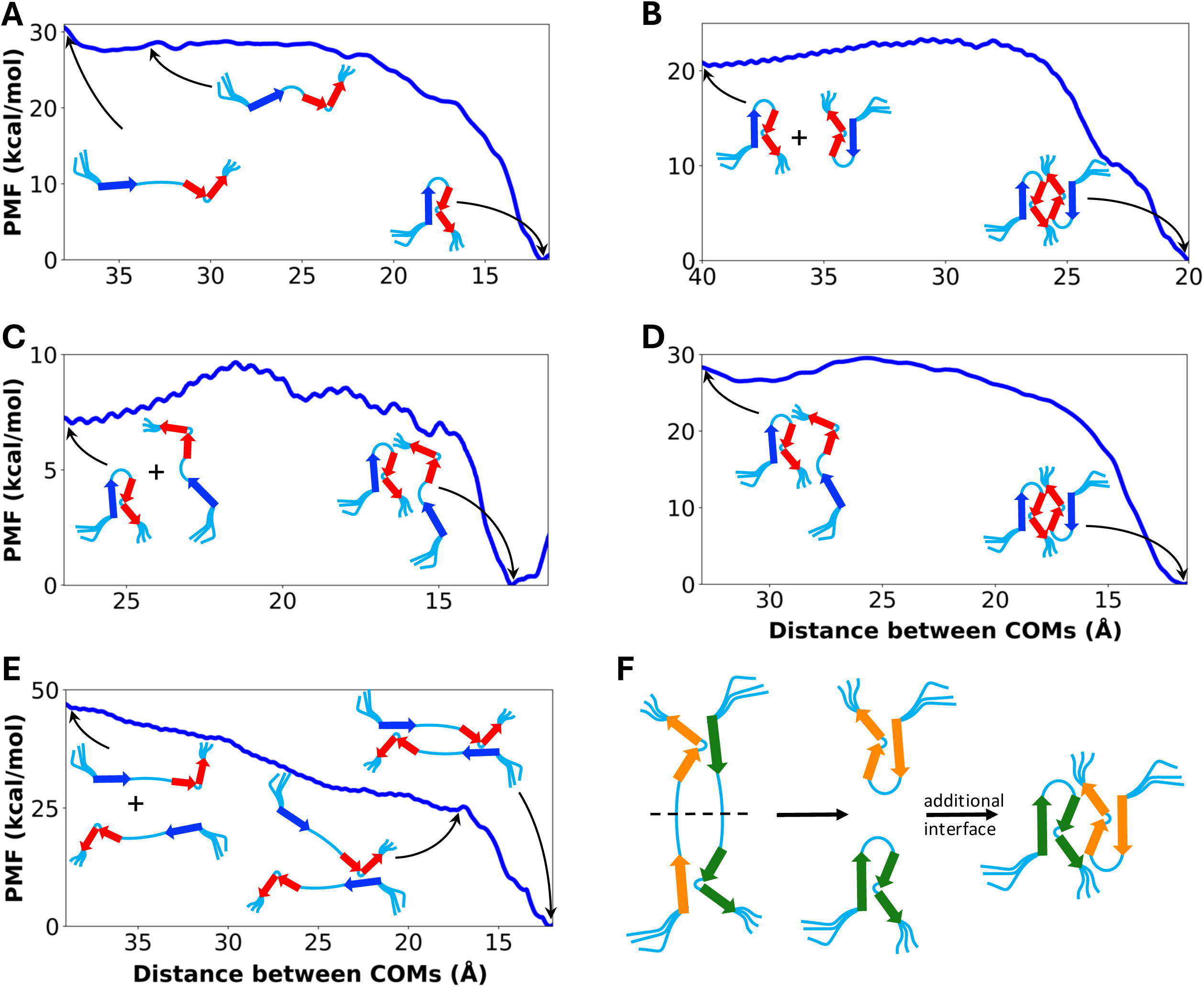
Free energy profiles for the evolution of the oligomeric seed in solution. (A) The free energy increases as a fibrillar subunit transitions from an R shape to semi-open to an open shape. (B) The free energy increases when the interface between the concave sides of two bent C-sheets is broken. Both subunits have an R shape. (C) Same as in (B) except one subunit now has a semi-open shape. (D) The free energy increases when an R-shaped subunit within a dimer unfolds into a semi-open shape. (E) The free energy increases as the two subunits in an open-shaped dimer are separated. An intermediate where only one of the two interfaces involving a straight N-sheet and a bent C-sheet is broken can be identified. N- and C-sheets are shown in blue and red, respectively. (F) The open-shaped dimeric protofilament as a domain-swapped version of the R-shaped dimeric protofilament.

We used targeted MD to change a tetramer in the R shape of the Kenyaga et al. ^18^ fibril toward a semi-open shape and then change the semi-open tetramer toward the open shape in PDB 8OVK^19^. Subsequently we used the distance between the COMs of the N- and C-sheets (N-sheet truncated to residues K16 to A21 due to terminal fraying) as the reaction coordinate and performed umbrella sampling. The resulting free energy profile is shown in Fig. 5A. Relative to the semi-open shape, the open shape is less stable by 3 kcal/mol but the R shape is considerably more stable, by 28 kcal/mol. The high stability of the R shape is due to the interface between the straight N-sheet and the convex side of the bent C-sheet, in particular with hydrophobic interactions between L17 and L34/V36 and between F19 and I32/L34 (Fig. S9A). Note that D23 and K28 form an intra-chain salt bridge in the open shape but fan out into the solution in the R shape (Fig. S9). In the latter case, D23 or K28 or both residues line inter-subunit interfaces in some fibrils (Fig. S2B, D, E) ^33, 34^.

An R-shaped dimeric protofilament can be formed in two ways. The first is the direct binding of two R-shaped subunits. Dimers of different interfaces have been solved, often involving at least a part of the bent C-sheet (Fig. S2A, B, C, E) ^18, 33–35^. Here we limit to the dimer where the interface is formed entirely by the concave side of the bent C-sheet in each subunit ^18^. Using the distance between the COMs of the two subunits as the reaction coordinate, a free energy decrease of 21 kcal/mol upon forming this interface is found by umbrella sampling (Fig. 5B).

A second way to form an R-shaped dimer starts with one R-shaped subunit and one semi-open subunit. In the first step, the two subunits form an intermediate with an interface between the respective bent C-sheets. In this case, the free energy decrease from umbrella sampling is 7 kcal/mol (Fig. 5C). The latter value should be comparable to that found in Fig. 5B; the discrepancy can be attributed to the difference between the two reaction coordinates: though both are defined as the distance between the subunit COMs, they are still different because the shape of the second subunit differs between Figs. 5B and 5C (R vs. semi-open). We take the average of these two values, 14 kcal/mol, as the free energy decrease upon forming an interface between two bent C-sheets. In the second step, the N-sheet of the semi-open subunit in the intermediate folds back to the C-sheet (Fig. 5D). This step involves the formation of the interface between the straight N-sheet and the convex side of the bent C-sheet, and the free energy decrease from umbrella sampling is 28 kcal/mol. This number is identical to the value found for an analogous step shown in Fig. 5A, with the difference being the absence of the other subunit.

Lastly we modeled the formation of the open-shaped dimeric protofilament, by the direct binding of two open-shaped subunits. This protofilament contains two inter-subunit interfaces, each formed between an N-sheet and a bent C-sheet (Fig. S9B). The free energy profile for this process is shown in Fig. 5E. An intermediate can be identified, where only one of the two inter-subunit interfaces is formed. The free energy decrease is 22 kcal/mol. Subsequently, the second interface forms, resulting in another free energy decrease of 25 kcal/mol.

From the umbrella sampling results, two observations can be made. First, the interface between a straight N-sheet and a bent C-sheet provides strong stabilization (∼25 kcal/mol), whether within the same subunit (as in the R-shaped protofilament) or between two subunits (as in the open-shaped protofilament). Other interfaces, such as that formed by the concave side of the bent C-sheet, also stabilize protofilaments, but to a lesser extent. This observation explains why all the R-shaped protofilaments have the same intra-subunit interface between a straight N-sheet and a bent C-sheet but a variety of additional inter-subunit interfaces (Fig. S2). Second, the interfaces between a straight N-sheet and a bent C-sheet are similar to each other in the R- and open-shaped protofilaments (Fig. S9). In particular, both involve F19 from the N-sheet and L34 and V36 from the C-sheet, though there is a registry shift in the interfaces of the two types of protofilaments. The resulting stabilization is also similar, ∼25 kcal/mol per interface. Indeed, the open-shaped dimeric protofilament may be seen as a domain-swapped version of an R-shaped dimeric protofilament (Fig. 5F). Cutting the linkers between the N- and C-sheets in two fibrillar subunits of the open-shaped protofilament and reconnecting the linkers between two neighboring subunits produce two isolated R-shaped fibrillar subunits. Forming an additional interface between the two subunits then results in the R-shaped dimeric protofilament.

### Membrane association initiates multiple pathways to fibrils

Combining the steps presented above, we obtain a complete picture of the membrane-assisted aggregation of Aβ40, involving multiple pathways that lead to polymorphic fibril structures (Fig. 6). Unstructured monomers bind to a membrane and, due to the reduced dimensionality, coalesce into partially formed β-sheets. Membrane defects, in part due to the embedding of the β-sheets, then facilitate the release of the β-sheets. The better-formed C-sheet comes into solution before the less-formed N-sheet, producing a semi-open shape for the oligomeric seed. Two residues, Y10 and K28, both help stabilizing the β-sheets in the membrane environment and their ordered release.

**Fig. 6.**
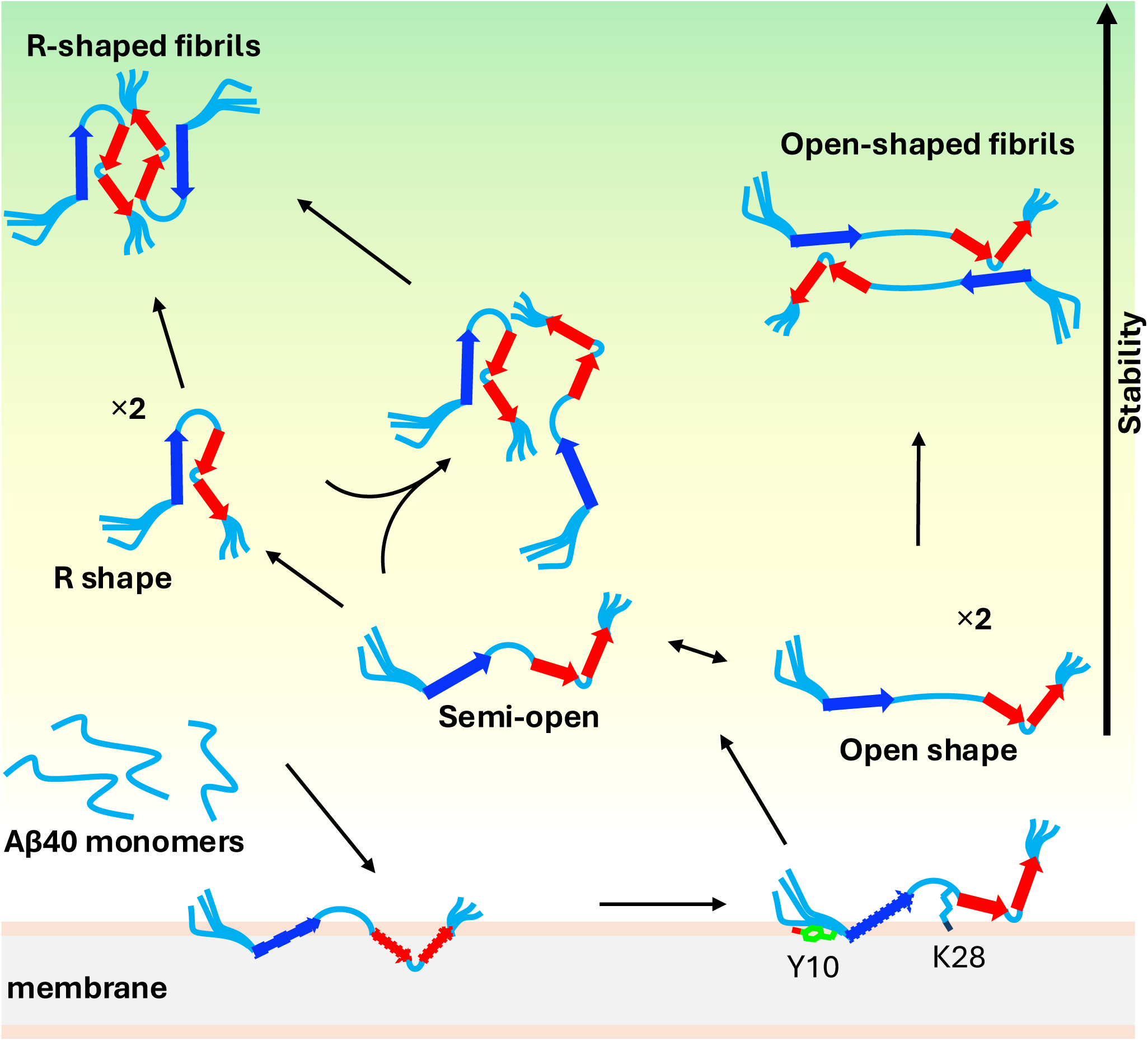
Membrane-assisted aggregation pathways of Aβ40. Unstructured Aβ40 monomers bind to the membrane and oligomerize to form β-sheets. By the time the C-sheet emerges in solution it is already well formed while the N-sheet is still incomplete and tethered to the membrane by Y10 and K28. Once released into solution, the oligomeric seed samples a range of shapes and dimerizes to form different protofilaments, ultimately leading to polymorphic fibrils.

The released oligomeric seed samples a range of shapes in solution. In one extreme, the straight N-sheet and the bent C-sheet fold into an R shape. In the opposite extreme, the two β-sheets separate from each other into an open shape. These subunits can further dimerize into stable protofilaments. Two R-shaped subunits can directly bind with each other to form a dimeric R-shaped protofilament. Alternatively, an R-shaped subunit and a semi-open subunit can dimerize into an intermediate, which becomes the dimeric R-shaped protofilament when the semi-open N-sheet folds back into the C-sheet in the same subunit. Lastly, two open-shaped subunits can directly bind with each other to form a dimeric open-shaped protofilament.

## Discussion

Using extensive MD simulations of Aβ40 tetramers in membranes as well as in solution, we have shown how the membrane environment accommodates and stabilizes a small oligomeric seed and how this seed, once released into solution, deforms and dimerizes to form different protofilaments. In the membranes, lipids facilely interact with the peptides. For example, by bending and by a small size, respectively, lipid acyl chains and cholesterol fill the space underneath the β-sheets. On the other hand, the tall headgroup of GM1 ganglioside can form multiple interactions with the elevated K28 residue. In solution, the interface between a straight N-sheet and a bent C-sheet provided major stabilization of protofilaments. Interestingly, this interface can form either within a single fibrillar subunit, or between two subunits, leading to two protofilaments related by a form of domain swap.

Our simulations present the first comprehensive, atomic-level picture of the multiple pathways for the membrane-assisted aggregation of Aβ40, and are supported by multiple lines of experimental evidence. Interrogation of intermediates in the lag phase of membrane-associated fibrillation by Kenyaga et al. ^18^ revealed the assembly and release of the C-sheet while the N-terminal residues are still membrane-bound. In our simulations, the C-sheet once formed remains intact but the N-sheet is prone to fraying. Moreover, according to steered MD, the C-sheet is released before the N-sheet, due to the tethering of Y10 but also K28. The tethering is in the forms of cation-π interaction and salt bridge, respectively, with lipid headgroups. In cryo-EM images of the most populated lipidic fibril obtained by Frieg et al. ^19^, both Y10 and K28 are surrounded by lipid densities. Binding assays of Aβ fragments identified K28 as an important residue for electrostatic interactions with anionic membranes ^16^. Two studies highlighted the role of Y10 in membrane tethering of channel-forming Aβ42 oligomers, by fluorescence spectroscopy ^9^ and by a combination of solution NMR and MD simulations ^55^.

Our work also reinforces other MD simulation studies of amyloidogenic proteins in membranes. For example, Y10 interactions with POPC and S26-K28 interactions with GM1 were observed in Aβ42 dimer simulations ^48^, and K28 interactions with POPC were observed in Aβ42 monomer simulations ^50^. Paul et al. ^52^ attributed the stability of FUS low-complexity fibrils to strengthened backbone hydrogen bonds at membrane surfaces relative to the aqueous environment. Indeed, backbone hydrogen bonds are thought to be even stronger in the hydrophobic core of the membrane due to its low dielectric constant ^56^. As a result, a correlation between burial depth and backbone hydrogen bonding strength is expected and is precisely what is exhibited by the isolated N- and C-sheets (Fig. S6). More broadly, our study of Aβ40 tetramers may serve as a model for characterizing the membrane-assisted aggregation pathways of other amyloidogenic proteins by MD simulations.

The importance of membranes may explain why fibrils from *in vitro* preparations without lipids sometimes fail to replicate those derived from patients. In the case of Aβ40, all patient-derived and membrane-exposed fibrils share a common feature, i.e., a straight N-sheet but a bent C-sheet. Furthermore, all membrane-exposed fibrils resemble at least one patient-derived fibril. For example, the membrane-exposed fibril structure 8OVK ^19^, with an open-shaped subunit, shows remarkable similarity to the sporadic AD patient-derived fibril structure 6W0O ^28^. Likewise, the membrane-exposed fibril structures of Niu et al. ^17^ and Kenyaga et al. ^18^ and three patient-derived fibril structures ^33, 34^ have a common R-shaped subunit (Fig. S2). In contrast, eight lipid-free *in vitro* fibril structures have a straight C-sheet that is folded back onto the N-sheet ^22–26^; the resulting U-shaped subunit has not been seen in any patient-derived fibril. Thus lipids may play crucial roles in determining fibril structures and be involved in endogenous fibrillation.

The facile interactions with lipids demonstrated here for Aβ40 tetramers suggest that small oligomers of amyloidogenic proteins can easily take advantage of the environment presented to them to accelerate an otherwise extremely slow nucleation process. Moreover, the fibril structures eventually formed may be dictated by aggregation pathways instead of absolute stability, as different pathways may be separated by high free energy barriers. This reasoning explains both the polymorphism and the disease specificity of the fibrils formed by the same amyloidogenic protein, as different disease conditions may present different environments (e.g., lipid composition) to the protein and each environment may favor a particular aggregation pathway. Aggregation pathways and their influence by environmental factors such as cell membranes should thus be studied as an integral part of disease mechanisms. Also importantly, an intermediate in a pathway may be destabilized or its progression to the next intermediate may be blocked. Pathways may thus present unique opportunities for developing targeted therapies.

## Materials and Methods

### System preparation for MD simulations

Peptide-membrane systems were prepared by using the CHARMM-GUI web server ^57^. Initially, the membrane was symmetric, each with 200 or 100 lipids per leaflet and a composition mimicking rat synaptic plasma membranes. Specifically, each leaflet contained 92, 50, 24, 14, 10, 6, and 4 copies of POPC, cholesterol, PSM, POPE, POPI, GM1, and POPS, respectively, for constructs containing both N- and C-sheets; the lipid numbers were halved for constructs with a single β- sheet. After including Aβ40 peptides, additional POPC lipids were added to the opposite leaflet to ensure equal surface areas in the two leaflets.

For the simulations of a single tetrameric β-sheet, the initial structures of the N- and C-sheets (residues Q15-D23 and K28-G38, respectively) were taken from PDB 6W0O ^28^. Each β-sheet was placed parallel to the membrane at four different depths (Fig. S3A, B). For the oligomeric seed, the initial structure was generated using the converged structures of the N- and C-sheets from the preceding step as templates, with disordered residues including Y10-H14, V24-N27, and V39-V40 built using MODELLER ^58^.

For all the simulations in solution, constructs composed of chains comprising residues Q15-G38 were used and solvated in a cubic box. Na^+^ and Cl^-^ ions were added to the systems either in membranes or in solution for charge neutralization and to maintain the salt concentration at 150 mM.

### MD Simulations

MD simulations were run in NAMD 3.0 ^59^ (with the exception described below) using the CHARMM36 ^60^ force field for proteins and lipid bilayers and the TIP3P ^61^ model for water. For each system, the energy was minimized for 10000 steepest-descent cycles, followed by a six- or three-step equilibration, for the systems in membranes and in solution, respectively. For the systems in membranes, the first two steps were at constant temperature and volume and the remaining four were at constant temperature and pressure. In each of the first three steps, the timestep was 1 fs and the simulation time was 125 ps; in the last three steps, the timestep was 2 fs and the simulation time was 500 ps. Heavy atoms of the peptides were position-restrained, with the force constant reducing from 10 to 0.1 kcal/mol/Å^2^; headgroups of lipids were also position-restrained, with the force constant reducing from 100 to 0 kcal/mol/Å^2^. For simulations in solution, the first step of equilibration was at constant temperature and volume and the remaining two were at constant temperature and pressure. The first two steps had a timestep of 1 fs and simulation time of 250 ps; the third step had a timestep of 2 fs and simulation time of 500 ps. The force constant for restraining the peptide heavy atoms was gradually reduced from 1.0 to 0.1 kcal/mol/Å^2^.

Production runs were carried out at constant temperature and volume without restraints. Bond lengths containing hydrogen atoms were constrained using the SHAKE algorithm ^62^. The particle mesh Ewald method ^63^ with a nonbonded cutoff of 12 Å was used to treat long-range electrostatic interactions. The temperature was maintained at 300 K by the Langevin thermostat ^64^ with a damping constant of 1 ps^-1^. The pressure was maintained at 1 atm using the Langevin piston ^65^ with an oscillation period of 50 ps and a decay time of 25 ps. Snapshots were saved every 100 ps for analysis.

From the simulations of the oligomeric seed embedded in membranes, a frame at 152.26 ns in one of the four replicates was chosen for the steered MD simulations to affect the release into solution. The COM of all Cα atoms was pulled at a speed of 1 Å/ns directly away from the membrane, and simulations were carried out in four replicates. A semi-open structure at 15.2 ns in one of the replicates was used to prepare for the umbrella sampling simulations described next. Details of all the simulations in membranes are summarized in Supplementary Table 1.

### Umbrella sampling

Umbrella sampling consisted of two parts. The first was to generate initial structures within different windows along a reaction coordinate; the second was to run simulations within each window, with the reaction coordinate restrained to the center of the window. We carried out the first part by running either targeted or steered MD simulations (Supplementary Table 2). The windows were separated by 0.25 Å for targeted MD and by 0.5 Å for steered MD.

Before targeted or steered MD simulations, the starting and target structures were further equilibrated for 10 ns after the brief equilibration in solution described above. Further equilibration was applied to the semi-open structure selected from the steered MD simulations in membranes; the R-shaped dimeric fibril structure of Kenyaga et al. ^18^, after uniformization by replicating a single chain eight times; and the open-shaped lipidic fibril structure 8OVK ^19^. Hydrogen bonds in the C- and N-sheets were restrained in the first 8 ns with a force constant of 10 kcal/mol/Å^2^; the restraints were released in the last 2 ns.

The equilibrated semi-open structure was used as the starting structure for the targeted MD simulations in the 33 Å to 38 Å portion of Fig. 5A and the target structure for the targeted MD simulations of Fig. 5D. For the 11.5 Å to 33 Å portion of Fig. 5A, the structures for starting umbrella sampling simulations were the same as those of the second subunit for Fig. 5D. The equilibrated R-shaped structure (or subunit thereof) was used as the starting structure for targeted MD simulations in the 11.5 Å to 33 Å portion of Fig. 5A, for targeted MD simulations of Fig. 5D, and for steered MD simulations of Fig. 5B. The equilibrated open-shaped structure was used as the target structure for the targeted MD simulations in the 33 Å to 38 Å portion of Fig. 5A and as the starting structure for the steered MD simulations of Fig. 5E. The final structure from the targeted MD simulations of Fig. 5D was used as the starting structure for the steered MD simulations of Fig. 5C.

Steered MD was used to separate the two subunits in a dimeric protofilament. The COM of the first subunit was fixed while the COM of the second subunit was pulled at a speed of 1 Å/ns along the initial inter-COM vector. For this part, COMs were calculated using backbone heavy atoms. Targeted MD was used to change the conformation of an isolated fibrillar subunit or a subunit within a dimeric protofilament. A root-mean-square-deviation (RMSD) restraint was imposed on the entire subunit, with a force constant of 500 kcal/mol/Å^2^. The RMSD was calculated between the current structure and the target structure using backbone heavy atoms.

For both steered and target MD, the reaction coordinate, defined as the distance between COMs of selected Cα atoms listed in Supplementary Table 2, was calculated. Frames where the reaction coordinate had values closest to the centers of the individual windows were saved for starting the second part of umbrella sampling, which was run in AMBER22 ^66^ on GPUs using *pmemd.cuda* ^67^.

Within each window, the simulation was run for 10 ns, with the reaction coordinate restrained to the center with a force constant of 10 kcal/mol/Å^2^. The weighted histogram analysis method ^68^ was applied to the data from all the windows to calculate the free energy profile.

### MD data analysis

A contact was defined when any heavy atom of a residue was within 3.5 Å of any heavy atom of the membrane. The membrane contact probability for each residue was determined by the proportion of frames in which that residue made at least one contact with the membrane. Ztip distances were calculated using a custom Python code by measuring the difference between the Z coordinate of the heavy tip atom of each sidechain and the average Z coordinate of phosphorous atoms in the leaflet embedding the Aβ40 peptides. The twist angles of β-sheets, defined as between inter-terminus vectors of successive β-strands ^54^, were calculated using a custom tcl code in VMD^69^. The first (second) end of a β-strand was identified as the midpoint between the carbonyl carbon of the first (second last) residue and the amide nitrogen of the second (last) residue. For the N-sheet, the N-terminal residues were 16 and 17 and the C-terminal residues were 20 and 21. For the bent C-sheet, a twist angle was calculated for each half, with terminal residues 29, 30, 32, and 33 for the first half and 33, 34, 36, and 37 for the second half. The number of hydrogen bonds was calculated using the *hbond* plugin in CPPTRAJ ^70^, with the distance and angle cutoffs at 3.5 Å and 125°, respectively. Line plots were made in Python 3.9.7, with smoothing by a Savitzky-Golay filter ^71^ from the SciPy library.

## Acknowledgement

We thank Dr. Wei Qiang for sharing the coordinates of their fibril structure. This work was supported by National Institutes of Health Grant AG073434.

## References

1. Vassar, R., et al. β-Secretase cleavage of Alzheimer’s amyloid precursor protein by the transmembrane aspartic protease BACE. Science 286, 735–741 (1999).

2. Wolfe, M. S., et al. Peptidomimetic probes and molecular modeling suggest that Alzheimer’s γ-secretase is an intramembrane-cleaving aspartyl protease. Biochemistry 38, 4720–4727 (1999).

3. Kukar, T. L., et al. Lysine 624 of the amyloid precursor protein (APP) is a critical determinant of amyloid beta peptide length: support for a sequential model of gamma-secretase intramembrane proteolysis and regulation by the amyloid beta precursor protein (APP) juxtamembrane region. J Biol Chem 286, 39804–39812 (2011).

4. Kang, J., et al. The precursor of Alzheimer’s disease amyloid A4 protein resembles a cell-surface receptor. Nature 325, 733–736 (1987).

5. Wolfe, M. S. In search of pathogenic amyloid β-peptide in familial Alzheimer’s disease. Prog Mol Biol Transl Sci 168, 71–78 (2019).

6. Hardy, J., Salih, D. TREM2-mediated activation of microglia breaks link between amyloid and tau. Lancet Neurol 20, 416–417 (2021).

7. Sarkar, B., Das, A. K., Maiti, S. Thermodynamically stable amyloid-β monomers have much lower membrane affinity than the small oligomers. Front Physiol 4, 47572 (2013).

8. Bode, D. C., Baker, M. D., Viles, J. H. Ion Channel Formation by Amyloid-beta42 Oligomers but Not Amyloid-beta40 in Cellular Membranes. J Biol Chem 292, 1404–1413 (2017).

9. Karkisaval, A. G., et al. The structure of tyrosine-10 favors ionic conductance of Alzheimer’s disease-associated full-length amyloid-beta channels. Nat Commun 15, 1296 (2024).

10. Tian, Y., Liang, R., Kumar, A., Szwedziak, P., Viles, J. H. 3D-visualization of amyloid-β oligomer interactions with lipid membranes by cryo-electron tomography. Chem Sci 12, 6896–6907 (2021).

11. Bode, D. C., Freeley, M., Nield, J., Palma, M., Viles, J. H. Amyloid-beta oligomers have a profound detergent-like effect on lipid membrane bilayers, imaged by atomic force and electron microscopy. J Biol Chem 294, 7566–7572 (2019).

12. Qiang, W., Kengewerere, M. K., Kenyaga, J. M. Modulation of Lipid Dynamics in the β- Amyloid Aggregates Induced Membrane Fragmentation. J Phys Chem B 128, 5667–5675 (2024).

13. Rosenberry, T. L., Zhou, H. X., Stagg, S. M., Paravastu, A. K. Oligomer Formation by Amyloid-β42 in a Membrane-Mimicking Environment in Alzheimer’s Disease. Molecules 27, (2022).

14. Ikeda, K., Yamaguchi, T., Fukunaga, S., Hoshino, M., Matsuzaki, K. Mechanism of amyloid beta-protein aggregation mediated by GM1 ganglioside clusters. Biochemistry 50, 6433–6440 (2011).

15. Banerjee, S., Hashemi, M., Zagorski, K., Lyubchenko, Y. L. Cholesterol in Membranes Facilitates Aggregation of Amyloid β Protein at Physiologically Relevant Concentrations. ACS Chem Neurosci 12, 506–516 (2021).

16. Chauhan, A., Ray, I., Chauhan, V. P. Interaction of amyloid beta-protein with anionic phospholipids: possible involvement of Lys28 and C-terminus aliphatic amino acids. Neurochem Res 25, 423–429 (2000).

17. Niu, Z., Zhao, W., Zhang, Z., Xiao, F., Tang, X., Yang, J. The molecular structure of Alzheimer β-amyloid fibrils formed in the presence of phospholipid vesicles. Angew Chem Int Ed Engl 53, 9294–9297 (2014).

18. Kenyaga, J. M., Cheng, Q., Qiang, W. Early stage β-amyloid-membrane interactions modulate lipid dynamics and influence structural interfaces and fibrillation. J Biol Chem 298, 102491 (2022).

19. Frieg, B., et al. Cryo-EM structures of lipidic fibrils of amyloid-β (1-40). Nat Commun 15, 1297 (2024).

20. Muhammedkutty, F. N. K., et al. A common pathway for detergent-assisted oligomerization of Aβ42. Commun Biol 6, 1184 (2023).

21. 21. Gao, Y., et al. Structural Model for Self-Limiting β-strand Arrangement Within an Alzheimer’s Amyloid-β Oligomer. bioRxiv, (2022).

22. Paravastu, A. K., Leapman, R. D., Yau, W. M., Tycko, R. Molecular structural basis for polymorphism in Alzheimer’s β-amyloid fibrils. Proc Natl Acad Sci U S A 105, 18349–18354 (2008).

23. Qiang, W., Yau, W. M., Luo, Y., Mattson, M. P., Tycko, R. Antiparallel β-sheet architecture in Iowa-mutant β-amyloid fibrils. Proc Natl Acad Sci U S A 109, 4443–4448 (2012).

24. Sgourakis, N. G., Yau, W. M., Qiang, W. Modeling an in-register, parallel “iowa” Aβ fibril structure using solid-state NMR data from labeled samples with Rosetta. Structure 23, 216–227 (2015).

25. Hu, Z. W., Vugmeyster, L., Au, D. F., Ostrovsky, D., Sun, Y., Qiang, W. Molecular structure of an N-terminal phosphorylated beta-amyloid fibril. Proc Natl Acad Sci U S A 116, 11253–11258 (2019).

26. Cerofolini, L., et al. Mixing Abeta(1-40) and Abeta(1-42) peptides generates unique amyloid fibrils. Chem Commun (Camb) 56, 8830–8833 (2020).

27. Kollmer, M., et al. Cryo-EM structure and polymorphism of Abeta amyloid fibrils purified from Alzheimer’s brain tissue. Nat Commun 10, 4760 (2019).

28. Ghosh, U., Thurber, K. R., Yau, W. M., Tycko, R. Molecular structure of a prevalent amyloid-β fibril polymorph from Alzheimer’s disease brain tissue. Proc Natl Acad Sci U S A 118, (2021).

29. Yang, Y., et al. Cryo-EM structures of Aβ40 filaments from the leptomeninges of individuals with Alzheimer’s disease and cerebral amyloid angiopathy. Acta Neuropathol Commun 11, 191 (2023).

30. Fu, Z., et al. An electrostatic cluster guides Abeta40 fibril formation in sporadic and Dutch-type cerebral amyloid angiopathy. J Struct Biol 216, 108092 (2024).

31. Pfeiffer, P. B., Ugrina, M., Schwierz, N., Sigurdson, C. J., Schmidt, M., Fandrich, M. Cryo-EM Analysis of the Effect of Seeding with Brain-derived Aβ Amyloid Fibrils. J Mol Biol 436, 168422 (2024).

32. Lu, J. X., Qiang, W., Yau, W. M., Schwieters, C. D., Meredith, S. C., Tycko, R. Molecular structure of β-amyloid fibrils in Alzheimer’s disease brain tissue. Cell 154, 1257–1268 (2013).

33. Fernandez, A., et al. Cryo-EM structures of amyloid-β and tau filaments in Down syndrome. Nat Struct Mol Biol 31, 903–909 (2024).

34. Yang, Y., et al. Cryo-EM structures of amyloid-β filaments with the Arctic mutation (E22G) from human and mouse brains. Acta Neuropathol 145, 325–333 (2023).

35. Schutz, A. K., et al. Atomic-resolution three-dimensional structure of amyloid β fibrils bearing the Osaka mutation. Angew Chem Int Ed Engl 54, 331–335 (2015).

36. Fitzpatrick, A. W. P., et al. Cryo-EM structures of tau filaments from Alzheimer’s disease. Nature 547, 185–190 (2017).

37. Falcon, B., et al. Tau filaments from multiple cases of sporadic and inherited Alzheimer’s disease adopt a common fold. Acta Neuropathol 136, 699–708 (2018).

38. Falcon, B., et al. Structures of filaments from Pick’s disease reveal a novel tau protein fold. Nature 561, 137–140 (2018).

39. Falcon, B., et al. Novel tau filament fold in chronic traumatic encephalopathy encloses hydrophobic molecules. Nature 568, 420–423 (2019).

40. Zhang, W., et al. Novel tau filament fold in corticobasal degeneration. Nature 580, 283–287 (2020).

41. Shi, Y., et al. Structure-based classification of tauopathies. Nature 598, 359–363 (2021).

42. Arseni, D., et al. Structure of pathological TDP-43 filaments from ALS with FTLD. Nature 601, 139–143 (2022).

43. Arseni, D., et al. TDP-43 forms amyloid filaments with a distinct fold in type A FTLD- TDP. Nature 620, 898–903 (2023).

44. Schweighauser, M., et al. Structures of α-synuclein filaments from multiple system atrophy. Nature 585, 464–469 (2020).

45. Yang, Y., et al. Structures of α-synuclein filaments from human brains with Lewy pathology. Nature 610, 791–795 (2022).

46. Yang, Y., et al. Cryo-EM structures of amyloid-β 42 filaments from human brains. Science 375, 167–172 (2022).

47. Brown, A. M., Bevan, D. R. Molecular Dynamics Simulations of Amyloid beta-Peptide (1-42): Tetramer Formation and Membrane Interactions. Biophys J 111, 937–949 (2016).

48. Fatafta, H., Khaled, M., Owen, M. C., Sayyed-Ahmad, A., Strodel, B. Amyloid-β peptide dimers undergo a random coil to beta-sheet transition in the aqueous phase but not at the neuronal membrane. Proc Natl Acad Sci U S A 118, (2021).

49. Nguyen, H. L., Linh, H. Q., Krupa, P., La Penna, G., Li, M. S. Amyloid β Dodecamer Disrupts the Neuronal Membrane More Strongly than the Mature Fibril: Understanding the Role of Oligomers in Neurotoxicity. J Phys Chem B 126, 3659–3672 (2022).

50. Wang, K., Shao, X., Cai, W. Binding Models of Aβ42 Peptide with Membranes Explored by Molecular Simulations. J Chem Inf Model 62, 6482–6493 (2022).

51. Christensen, M., Skeby, K. K., Schiott, B. Identification of Key Interactions in the Initial Self-Assembly of Amylin in a Membrane Environment. Biochemistry 56, 4884–4894 (2017).

52. Paul, S., Mondal, S., Shenogina, I., Cui, Q. The molecular basis for the increased stability of the FUS-LC fibril at the anionic membrane- and air-water interfaces. Chem Sci 15, 13788–13799 (2024).

53. Lomize, A. L., Todd, S. C., Pogozheva, I. D. Spatial arrangement of proteins in planar and curved membranes by PPM 3.0. Protein Sci 31, 209–220 (2022).

54. Ho, B. K., Curmi, P. M. Twist and shear in beta-sheets and beta-ribbons. J Mol Biol 317, 291–308 (2002).

55. Ciudad, S., et al. Aβ(1-42) tetramer and octamer structures reveal edge conductivity pores as a mechanism for membrane damage. Nat Commun 11, 3014 (2020).

56. Zhou, H. X., Cross, T. A. Influences of membrane mimetic environments on membrane protein structures. Annu Rev Biophys 42, 361–392 (2013).

57. Jo, S., Kim, T., Iyer, V. G., Im, W. CHARMM-GUI: a web-based graphical user interface for CHARMM. J Comput Chem 29, 1859–1865 (2008).

58. Webb, B., Sali, A. Comparative protein structure modeling using MODELLER. Curr Protoc Bioinformatics 54, 5.6.1–5.6.37 (2016).

59. Phillips, J. C., et al. Scalable molecular dynamics with NAMD. J Comput Chem 26, 1781–1802 (2005).

60. Klauda, J. B., et al. Update of the CHARMM all-atom additive force field for lipids: validation on six lipid types. J Phys Chem B 114, 7830–7843 (2010).

61. Jorgensen, W. L., Chandrasekhar, J., Madura, J. D., Impey, R. W., Klein, M. L. Comparison of Simple Potential Functions for Simulating Liquid Water. J Chem Phys 79, 926–935 (1983).

62. Ryckaert, J.-P., Ciccotti, G., Berendsen, H. J. C. Numerical integration of the cartesian equations of motion of a system with constraints: molecular dynamics of n-alkanes. Journal of Computational Physics 23, 327–341 (1977).

63. Essmann, U., Perera, L., Berkowitz, M. L., Darden, T., Lee, H., Pedersen, L. G. A Smooth Particle Mesh Ewald Method. J Chem Phys 103, 8577–8593 (1995).

64. Pastor, R. W., Brooks, B. R., Szabo, A. An Analysis of the Accuracy of Langevin and Molecular-Dynamics Algorithms. Mol Phys 65, 1409–1419 (1988).

65. Feller, S. E., Zhang, Y. H., Pastor, R. W., Brooks, B. R. Constant-Pressure Molecular-Dynamics Simulation - the Langevin Piston Method. J Chem Phys 103, 4613–4621 (1995).

66. 66. Case, D. A., et al. AMBER 2022, University of California, San Francisco. (ed^(eds) (2022).

67. Salomon-Ferrer, R., Gotz, A. W., Poole, D., Le Grand, S., Walker, R. C. Routine Microsecond Molecular Dynamics Simulations with AMBER on GPUs. 2. Explicit Solvent Particle Mesh Ewald. J Chem Theory Comput 9, 3878–3888 (2013).

68. Grossfield, A. WHAM: an implementation of the weighted histogram analysis method, Vesion 2.0.11. (ed^(eds). http://membrane.urmc.rochester.edu/content/wham.

69. Humphrey, W., Dalke, A., Schulten, K. VMD: visual molecular dynamics. J Mol Graph 14, 33–38, 27-38 (1996).

70. Roe, D. R., Cheatham, T. E., 3rd. PTRAJ and CPPTRAJ: Software for Processing and Analysis of Molecular Dynamics Trajectory Data. J Chem Theory Comput 9, 3084–3095 (2013).

71. Savitzky, A., Golay, M. J. E. Smoothing and Differentiation of Data by Simplified Least Squares Procedures. Anal Chem 36, 1627–1639 (1964).

