## Supplementary Tables and Figures for "Membrane-assisted Aβ40 aggregation pathways"

Supplementary Table 1. Details of simulations in membranes

| Residues | Simulation time (ns) | Box dimensions (Å$^3$) | # of atoms | # of replicates |
|---|---|---|---|---|
| Q15-D23 | 100[a] | 77 × 77 × 131 | 79756 | 1 |
| | | 77 × 77 × 127 | 78424 | 1 |
| | | 76 × 76 × 129 | 77635 | 1 |
| | | 73 × 73 × 129 | 71389 | 1 |
| K28-G38 | 100[b] | 78 × 78 Å × 128 | 80752 | 1 |
| | | 77 × 77 Å × 130 | 78803 | 1 |
| | | 75 × 75 Å × 130 | 75212 | 1 |
| | | 73 × 73 Å × 131 | 71936 | 1 |
| Y10-V40 | 600 | 108 × 108 × 182 | 217722 | 4 |
| Y10-V40 (steered MD) | 100 | 108 × 108 × 182 | 217722 | 4 |

[a]The N-sheet construct was placed at four depths relative to the membrane surface; the simulation was then carried out for 100 ns at each depth.

[b]The C-sheet construct was placed at four depths relative to the membrane surface; the simulation was then carried out for 100 ns at each depth.

Supplementary Table 2. Details of biased simulations in solution to generate initial structures for umbrella sampling

| Biasing method | Starting structure | Target structure | Box dimensions (Å$^3$) | # of atoms | Speed or force constant | # of windows | Residues for defining COMs |
|---|---|---|---|---|---|---|---|
| targeted | R shape | Semi-open | 87 × 87 × 87 | 66689 | 500 kcal/mol /Å$^2$ | 87 | K16-A21 / A30-V36 |
| targeted | Semi-open | Open shape | 88 × 88 × 88 | 68014 | 500 kcal/mol /Å$^2$ | 20 | K16-A21 / A30-V36 |
| steered | R-shaped dimer | separated subunits | 88 × 88 × 88 | 68121 | 1 Å/ns | 43 | 1$^{st}$ / 2$^{nd}$ subunits |
| steered | Open-shaped dimer | separated subunits | 103 × 103 × 103 | 109020 | 1 Å/ns | 56 | 1$^{st}$ / 2$^{nd}$ subunits |
| targeted | R-shaped dimer | R-shaped / semi-open dimer | 88 × 88 × 88 | 68121 | 500 kcal/mol /Å$^2$ | 87 | K16-A21 / A30-V36 of 2$^{nd}$ subunit |
| steered | R-shaped / semi-open dimer | separated subunits | 88 × 88 × 88 | 68121 | 1 Å/ns | 32 | 1$^{st}$ / 2$^{nd}$ subunits |

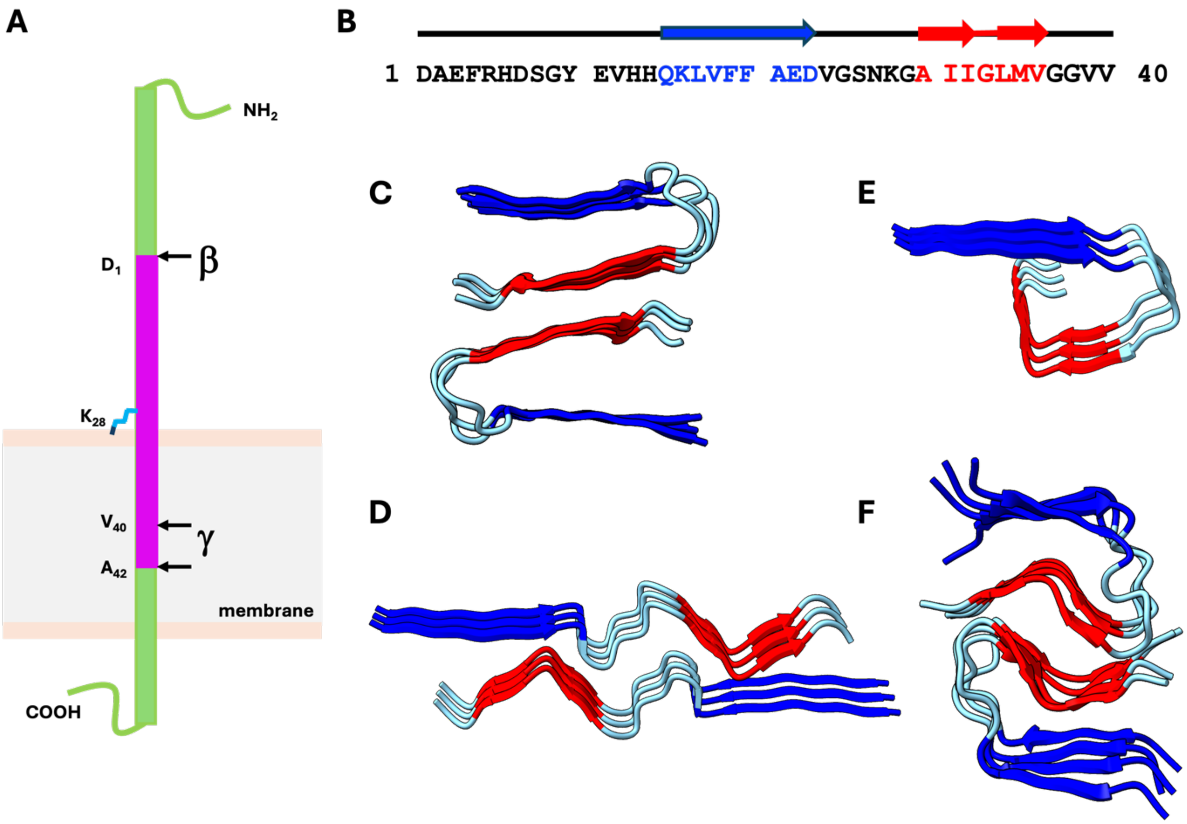


Fig. S1. Generation of Aβ peptides and sequence, secondary structure, and fibril structures of Aβ40. (A) Proteolytic cleavage of amyloid precursor protein. (B) Sequence and secondary structure of Aβ40. (C) U-shaped dimeric fibril in PDB 2LMN [22]. (D) Open-shaped dimer fibril in PDB 6W0O [28]. (E) Curved fibril in PDB 8OVM [19]. (F) R-shaped dimeric fibril of Kenyaga et al. [18]. The N- and C-sheets are shown in blue and red, respectively.

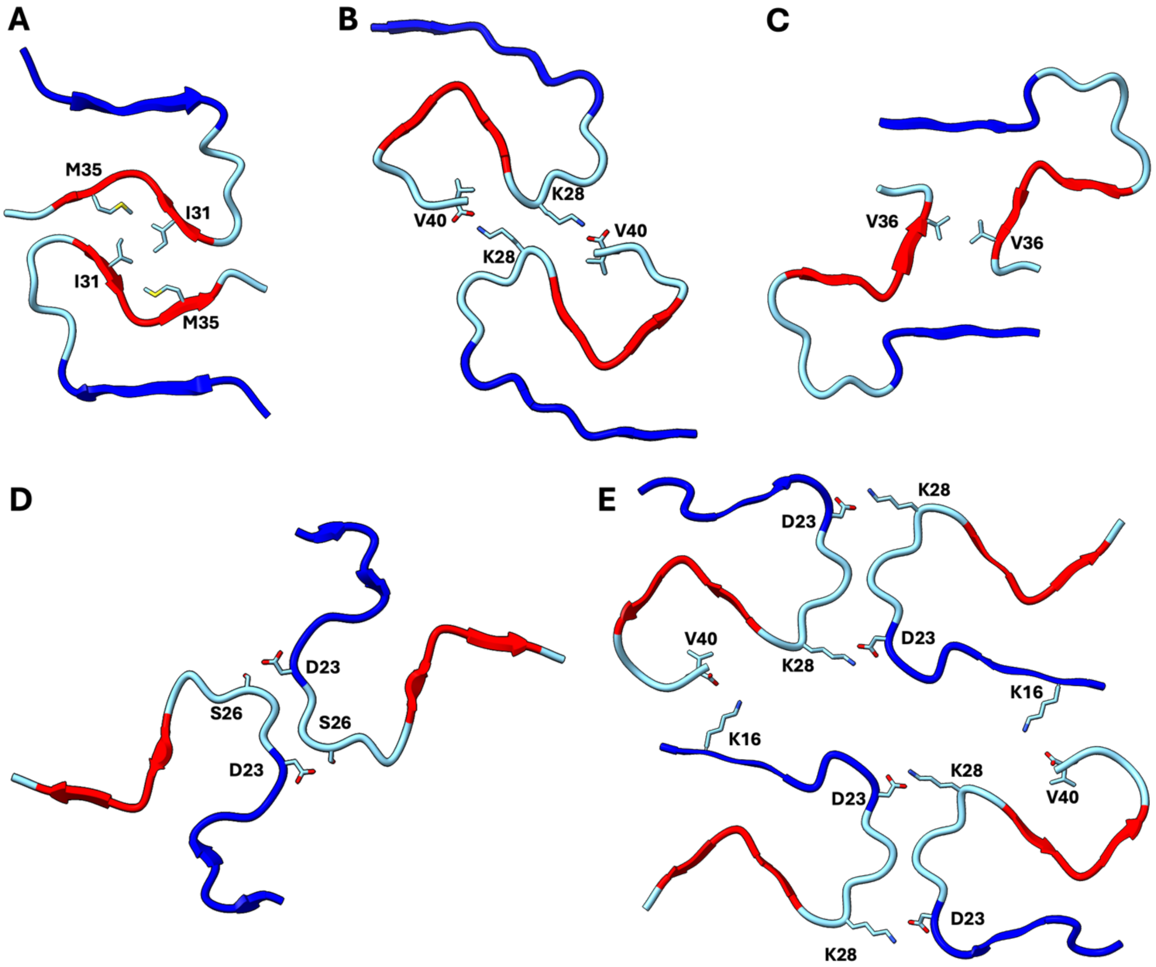


Fig S2. Dimers and tetramers of R-shaped protofilaments involve a variety of inter-subunit interfaces. (A) Dimeric fibril of Kenyaga et al. [18]. (B) Dimeric fibril in PDB 8SEK [33]. (C) Dimeric fibril in PDB 2MVX [35]. (D) Dimeric fibril in PDB 8BG9 [34]. (E) Tetrameric fibril in PDB 8BG0 [34]. N- and C-sheets are in blue and red, respectively. Some sidechains involved in the interfaces are shown as sticks.

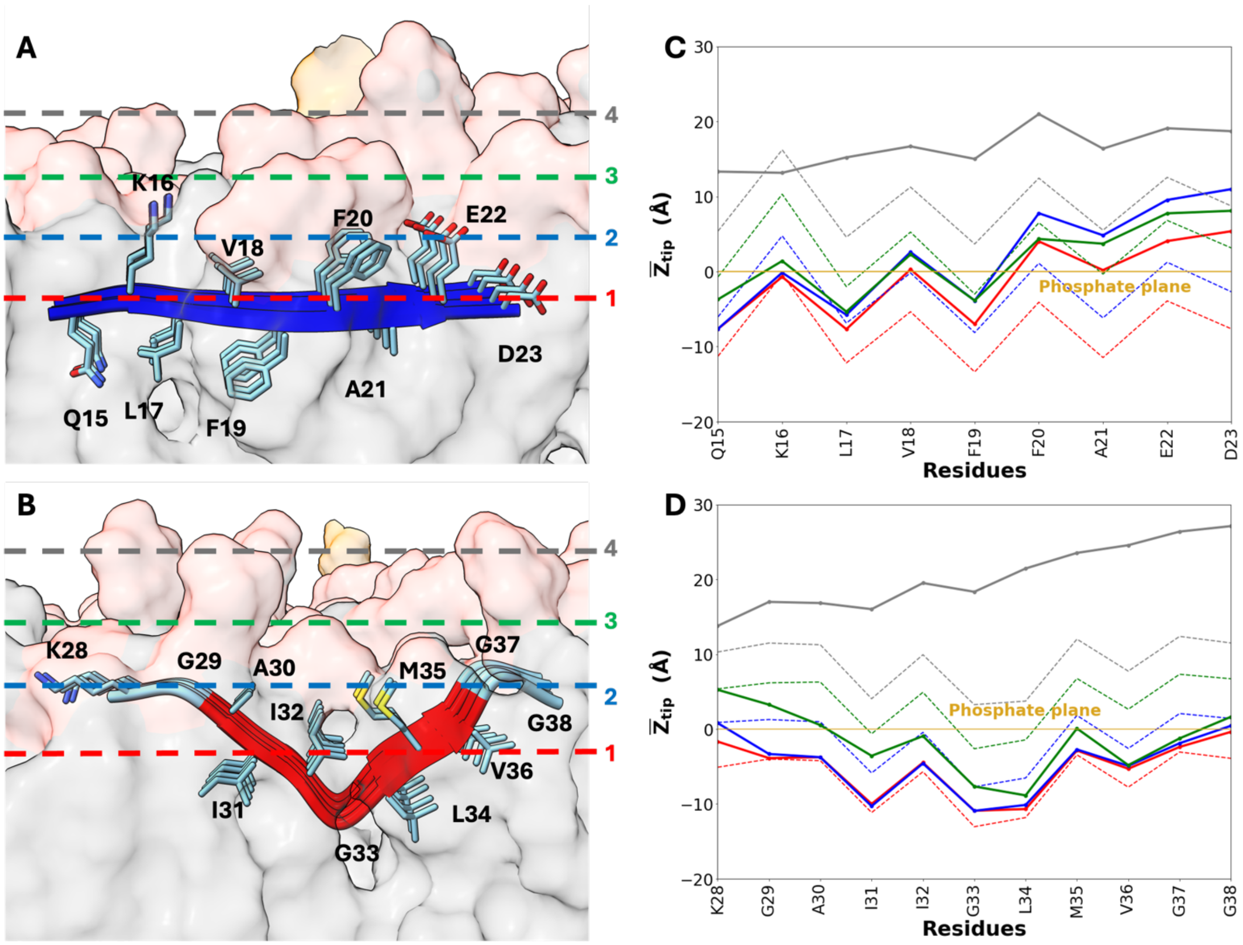


Fig. S3. Preferred burial depths of N- and C-sheets in membranes. (A, B) Same as Fig. 1A, B, except that four dashed lines are drawn to indicate the initial depth. The dashed lines are drawn at the average Z value of all heavy atoms; the structure shown is at depth 1. (C, D) The initial (dashed lines) and final (solid lines) mean $Z_{tip}$ distances for each residue in the N- and C-sheets. Results at the four depths are displayed in colors matching those of the dashed lines in panels (A) and (B). The final mean $Z_{tip}$ distances were averages over the last 10 ns of the 100-ns simulations.

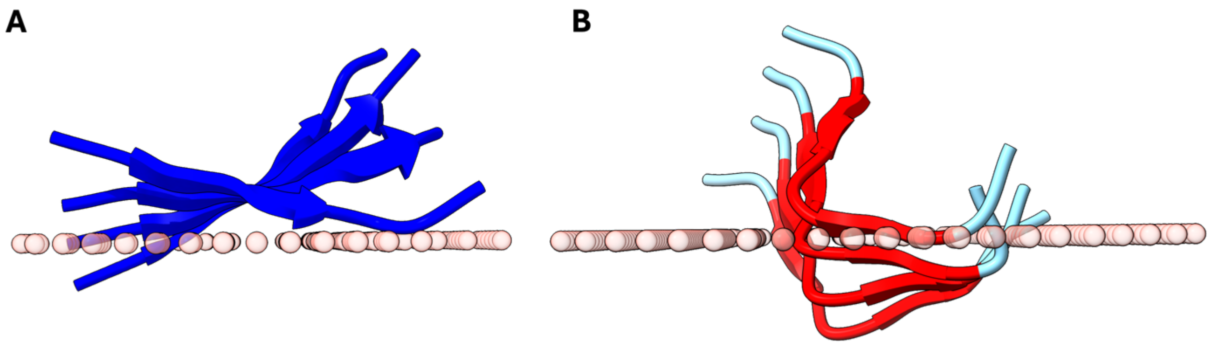


Fig. S4. Placement of the N- and C-sheets by the PPM web server. (A, B) N- and C-sheets. The phosphate plane is shown by pink spheres.

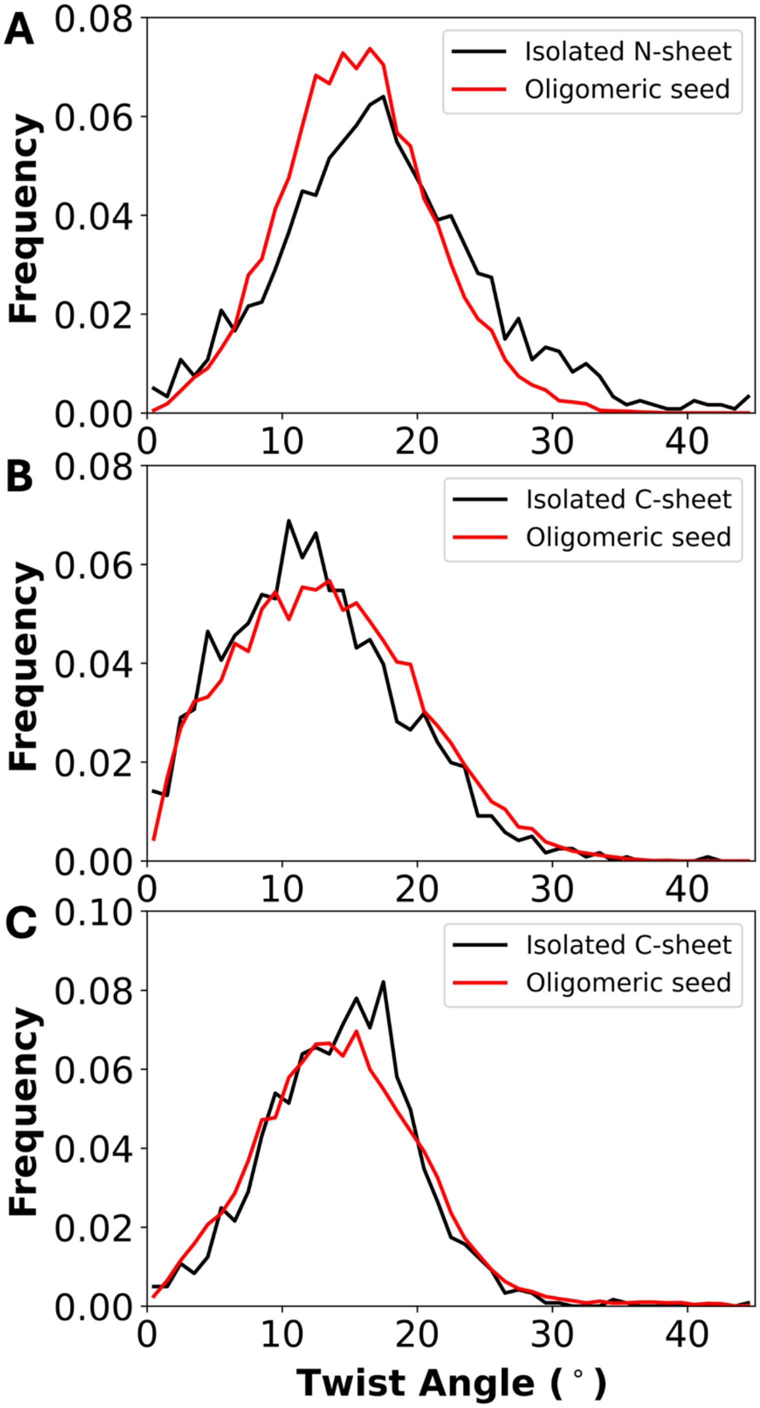


Fig. S5. Histograms of the twist angles of membrane-embedded N- and C-sheets. (A) N-sheet. (B) The first half of the C-sheet, before G33. (C) The second half of the C-sheet. In each frame, the twist angle of the β-sheet was the average of those calculated from three adjacent pairs of β-strands. For the isolated N- or C-sheet, the results from the simulations at burial depths 1 and 2 were averaged. For the oligomeric seed, results from four replicate simulations were averaged.

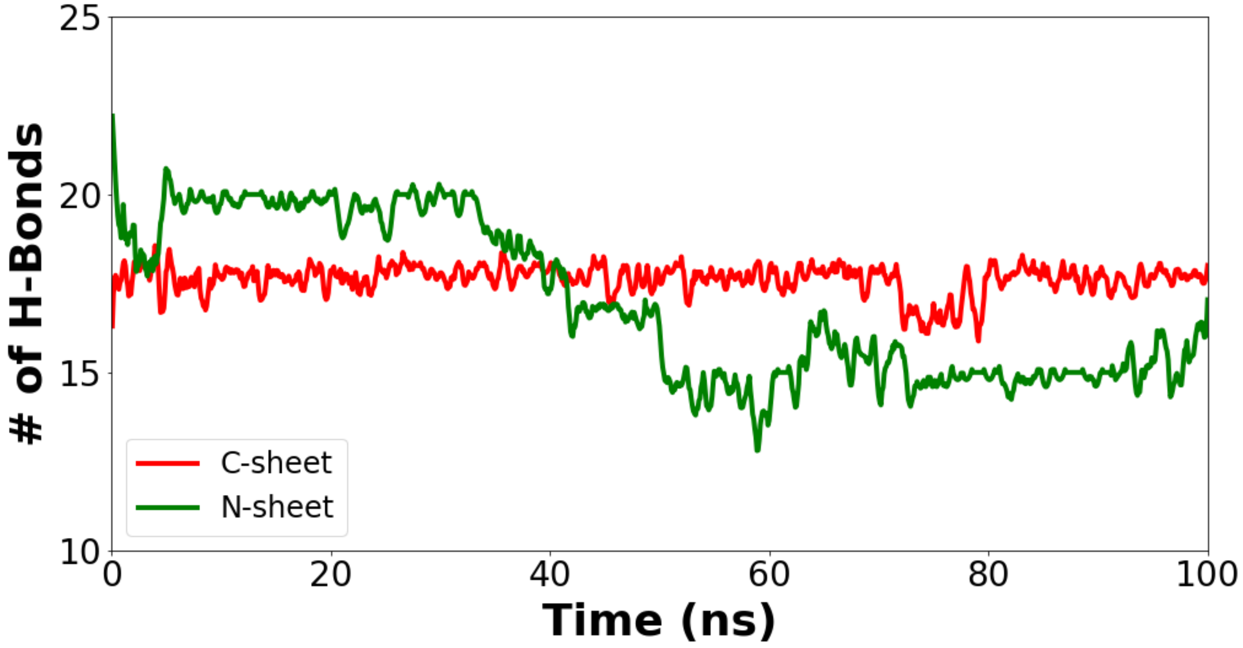

Fig. S6. Number of hydrogen bonds in the membrane-embedded N- or C-sheet. The results were from the simulations with the deepest initial burial.

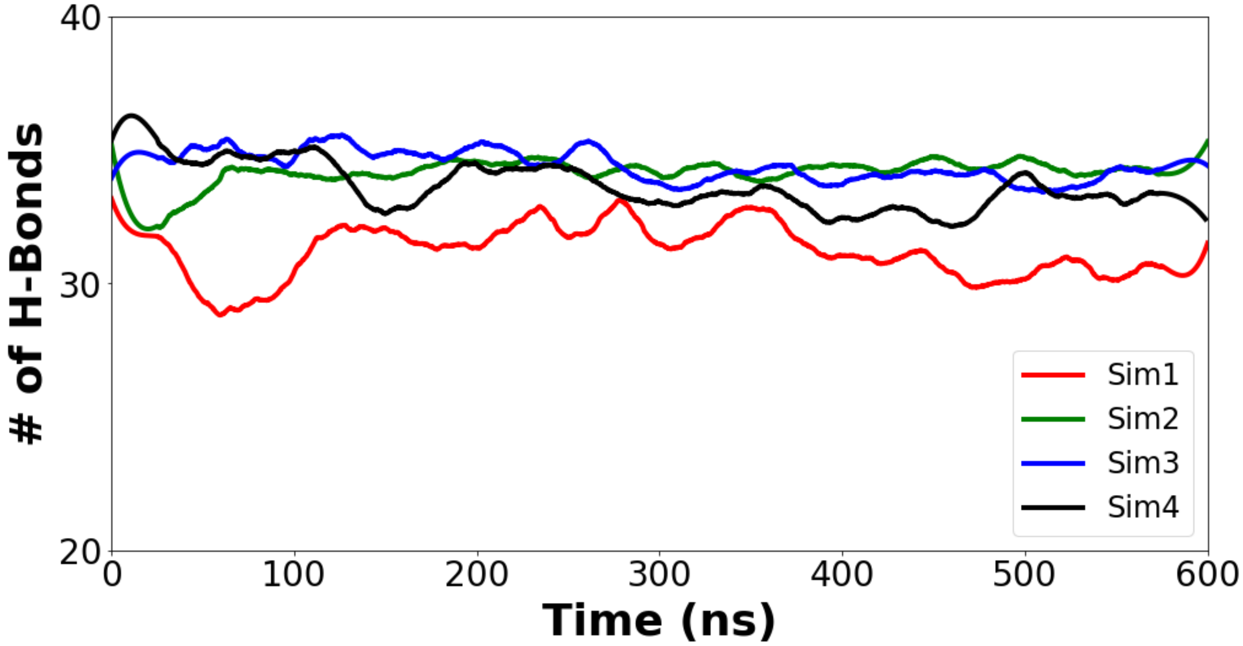

Fig. S7. Total number of hydrogen bonds in the membrane-embedded oligomeric seed.

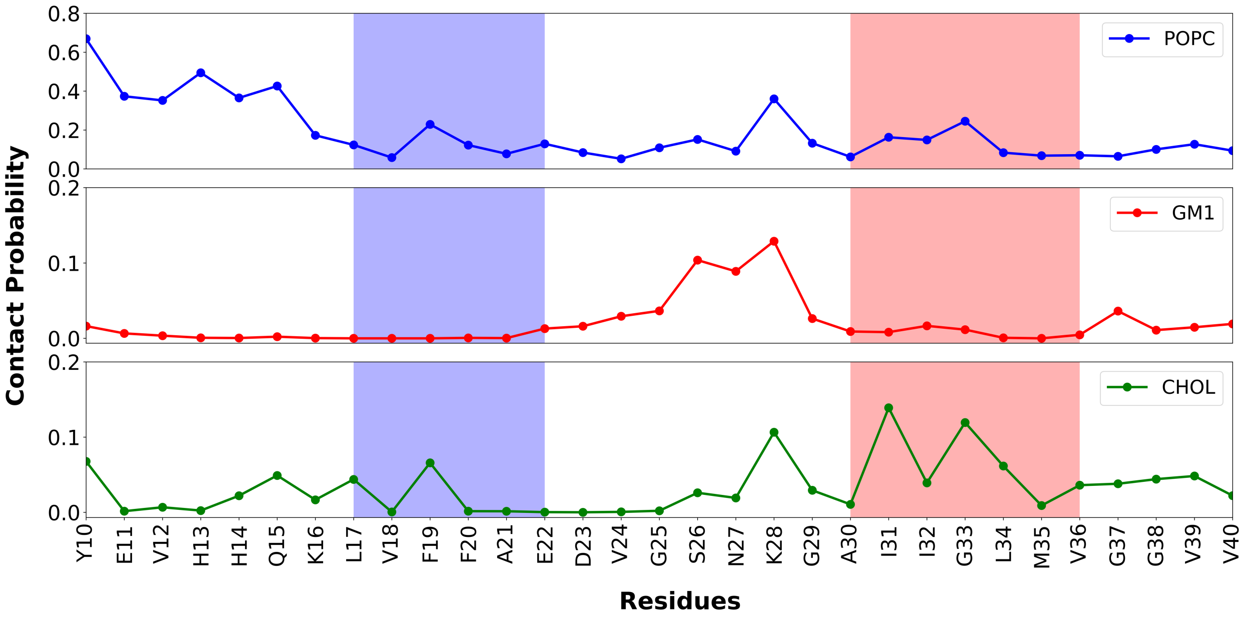


Fig. S8. Membrane contact probabilities of individual residues in the oligomeric seed, broken down according to lipid types. The results are similar to those in Fig. 3A, but each panel here reports contacts with a particular type of lipids in the membrane. The N- and C-sheets are indicated by blue and red shading, respectively.

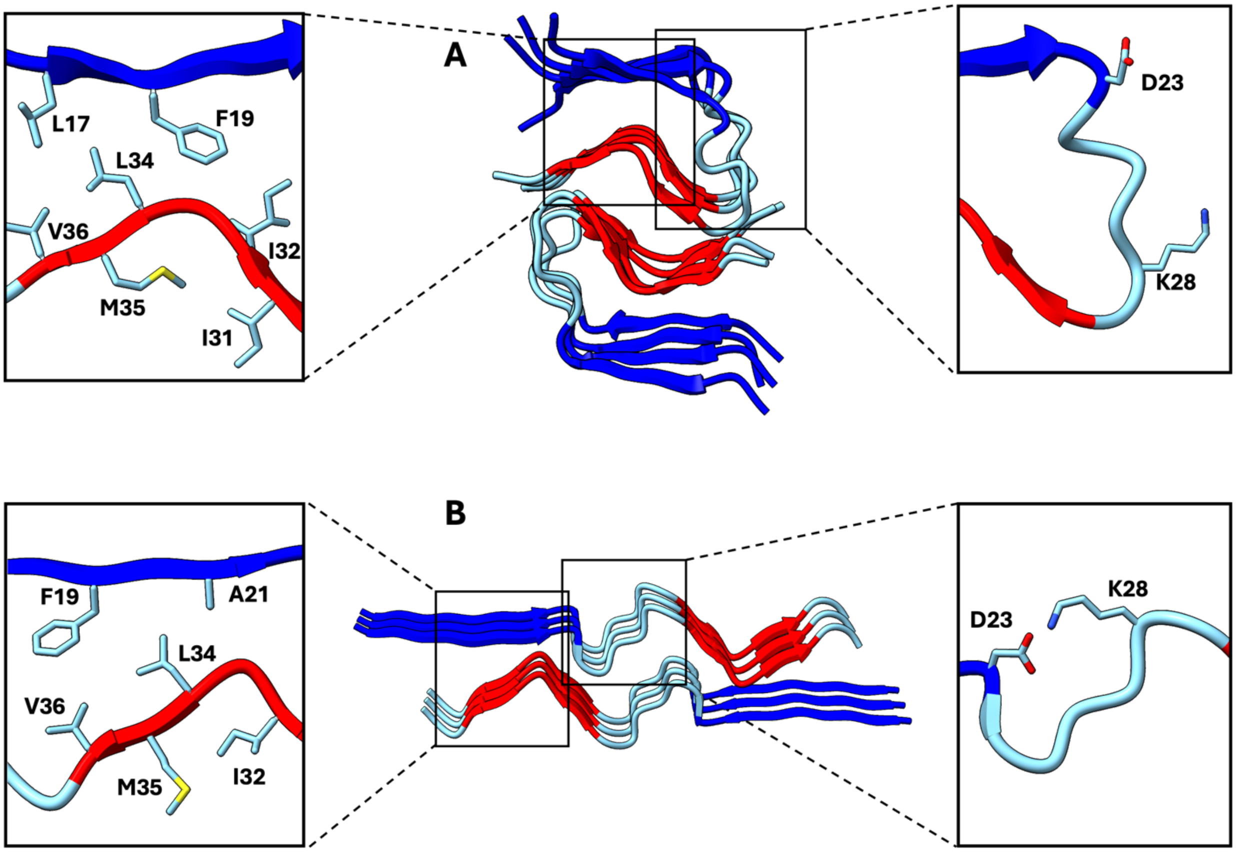


Fig. S9. Similarity in interfaces between the R- and open-shaped protofilaments. (A) R-shaped dimeric fibril of Kenyaga et al. [18]. (B) Open-shaped dimeric fibril in PDB 6W0O [28]. Zoomed views on the left highlight the interfaces between a straight N-sheet and a bent C-sheet, either within the same subunit or between two subunits. Zoomed views on the right show different arrangements of D23 and K28 in the two protofilaments.
